# How reliably can species co-occurrence be inferred from occupancy models?

**DOI:** 10.1101/2024.10.29.620848

**Authors:** Julie Louvrier, Mason Fidino, Josefa Vergara Stuardo, Olivier Gimenez

## Abstract

Interspecific interactions, such as predation, competition, and mutualism, are fundamental to the structure and function of any biotic community. Understanding these interactions provides valuable insights into community dynamics and informs conservation efforts. One common approach to studying these interactions is through co-occurrence patterns, which remain essential for understanding species relationships. Occupancy models have emerged as powerful tools to investigate these patterns, offering a flexible framework for ecological questions. Among these models, the co-occurrence framework developed by Rota et al. (2016) has seen widespread applications across various species and habitats.

In this study, we used simulations to evaluate the performance of the Rota et al. (2016) co-occurrence model under different study designs and species interaction scenarios. Specifically, we examined how the bias and precision of species occupancy estimates are influenced by the presence or absence of another species, varying the number of sampling units surveyed and the number of temporal replicates per unit. Additionally, we compared the co-occurrence model’s performance with a simpler model that does not account for species interactions to determine when statistically significant co-occurrence patterns emerge.

We observed that occupancy parameters become more precise as the intensity of species avoidance increased, and while the number of sampling sites significantly influenced precision, the number of temporal replicates did not. Notably, the co-occurrence model outperformed the non-interaction model only when species avoidance was strong and at least 150 sites were sampled. Based on these results, we recommend a minimum of 150 sampling sites when applying the Rota et al. (2016) model to evaluate co-occurrence between two potentially interacting species. Our study emphasizes the need for further development of methods capable of detecting interaction signals, even at low to moderate interaction levels, and in studies with fewer sampling sites.

## Introduction

Interspecific interactions such as predation, competition or mutualism represent a crucial component of any biotic community (Chesson, 2000; Letten et al., 2017). Such interactions are a major factor in niche segregation that can lower species demographic rates and restrain their spatial ranges (Linnell & Strand, 2000; Parsons et al., 2019). Understanding these interactions provides insights into community functioning and improves the conservation of communities (Singer et al., 2016).

To understand interspecific interactions, studies have often investigated co-occurrence patterns between species. However, assessing co-occurrence can be challenging due to the elusive nature of animal species that leads to false negatives on sites that are surveyed (MacKenzie et al., 2002; Mackenzie et al., 2003). As such, the occupancy modelling framework was developed to correct for the imperfect detection of species and other aspects of the data collection process (MacKenzie et al. 2002, Kéry et al. 2010; Kéry & Schaub 2011). Since their first introduction, occupancy models have been developed to address a wide array of ecological questions (see Bailey et al. 2013 for a review). For instance, occupancy models have been developed to estimate 1) co-occurrence patterns between a dominant and subordinate species (MacKenzie et al., 2017; Richmond et al., 2010; Waddle et al., 2010), 2) community-level dynamics (Dorazio et al., 2010; Dorazio & Royle, 2005; Kéry & Royle, 2009), and 3) interactions between two or more species within a single (MacKenzie et al., 2021; Rota et al., 2016) or multiple seasons (Fidino et al., 2019; Kleiven et al., 2023; Zipkin et al., 2023). As such, the comprehensive scope of occupancy models combined with their ability to address myriad ecological questions make them an indispensable tool in ecological research.

Of these extensions, the Rota et al. (2016) co-occurrence modelling framework has seen widespread use across a range of species and habitats (Fowler et al. 2021, Salvatori 2021, Twining et al. 2021, Saeed et al. 2022). As a generalization of the MacKenzie et al. (2002) co-occurrence modelling approach, the Rota et al. (2016) framework quantifies co-occurrence patterns by assuming the occupancy state at a site is a multivariate Bernoulli random variable. Yet, to quantify statistical associations between species no doubt requires a sufficient sample size of locations both with and without the species of interest. As a result, a key aspect of designing occupancy studies is to select an appropriate number of sampling units and temporal replicates per sampling unit, and many studies provide guidance on this topic for a variety of occupancy models (Guillera-Arroita & Lahoz-Monfort, 2012; MacKenzie et al., 2017; Mackenzie & Royle, 2005). Most of these studies, however, have focused on developing appropriate sampling designs to help ensure precise and unbiased occupancy estimates for a single species (Mackenzie & Royle, 2005). Given the widespread use of the Rota et al. (2016) framework, it is unfortunate that we lack sufficient guidelines to know whether varying study designs can produce reliable co-occurrence estimates, if a failure to detect statistical interactions is due to a lack of statistical power or know if model parameters can be estimated without too much bias and with satisfying precision.

Here, we used simulations to assess the performance of the Rota et al. (2016) co-occurrence modelling framework under varying study designs and interactions between species (DiRenzo et al., 2023). More specifically, we simulated the occurrence of two hypothetical species - species A is dominant and species B is subordinate - under patterns of increasing avoidance across varying sample sizes. To help guide future use of this model we investigated how the bias and precision of a species occupancy estimates given the presence or absence of the other species changes with the number of sampling units surveyed and the number of temporal replicates per unit. Furthermore, to understand when the model supports a statistically significant co-occurrence pattern, we compared the performance of this model to a model that did not account for species interactions. We hypothesized that the bias and the precision of co-occurrence parameter estimates would improve with (i) an increasing level of avoidance between the two species (i.e., effect size), but also (ii) with an increasing number of sites and temporal replicates. Secondly, we hypothesized that the model would support a significant interaction (iii) with no regards to the interaction level but (iv) the performance would increase with an increasing number of sites and temporal replicates.

## Methods

We adopted the model developed by Rota et al. (2016) in which we considered only 2 interacting species for simplicity, which we denote as species A and B. Following the formulation developed by Lonsinger (2022), we simulated the presence of B when influenced by the presence or absence of A with Ψ^A^ (the occupancy probability of A independent of B), Ψ^BA^ (the occupancy probability of B given the presence of A), and Ψ^Ba^ (the occupancy probability of B given the absence of A). We assumed a dominant-subordinate interaction in which A was the dominant species, and B was the subordinate one.

We used different occupancy scenarios for species A and B. For A, the dominant species, we tested two scenarios corresponding to Ψ^A^ = 0.5 and Ψ^A^ = 0.7. Species B, the subordinate, was distributed in seven scenarios with respect to A, which varied in how negatively B was associated with A (i.e. avoidance). We considered seven levels along a gradient of avoidance with an increasing difference between Ψ^BA^ and Ψ^Ba^. (Hereafter called “None”, “Low”, “Low-Medium”, “Medium”, “High-Medium”, “Low-High” and “High” see table 1 and discussion for interpretation of these levels). These scenarios depicted an interaction gradient from no avoidance (i.e. “None” scenario, in which Ψ^BA^ = Ψ^Ba^) to a strong avoidance of B by A (i.e. “High” scenario, in which Ψ^BA^ = 0.1 and Ψ^BA^ = 0.8). To set a baseline understanding of the study design required to fit the simplest model, we used a framework with no environmental variation (i.e. no covariate). To avoid perfect detectability in scenarios with a high number of temporal replicates, we used the probability of detecting a species at least once (θ) to calculate the detection probability of a single temporal replicate (*p*):

**Table 1:**
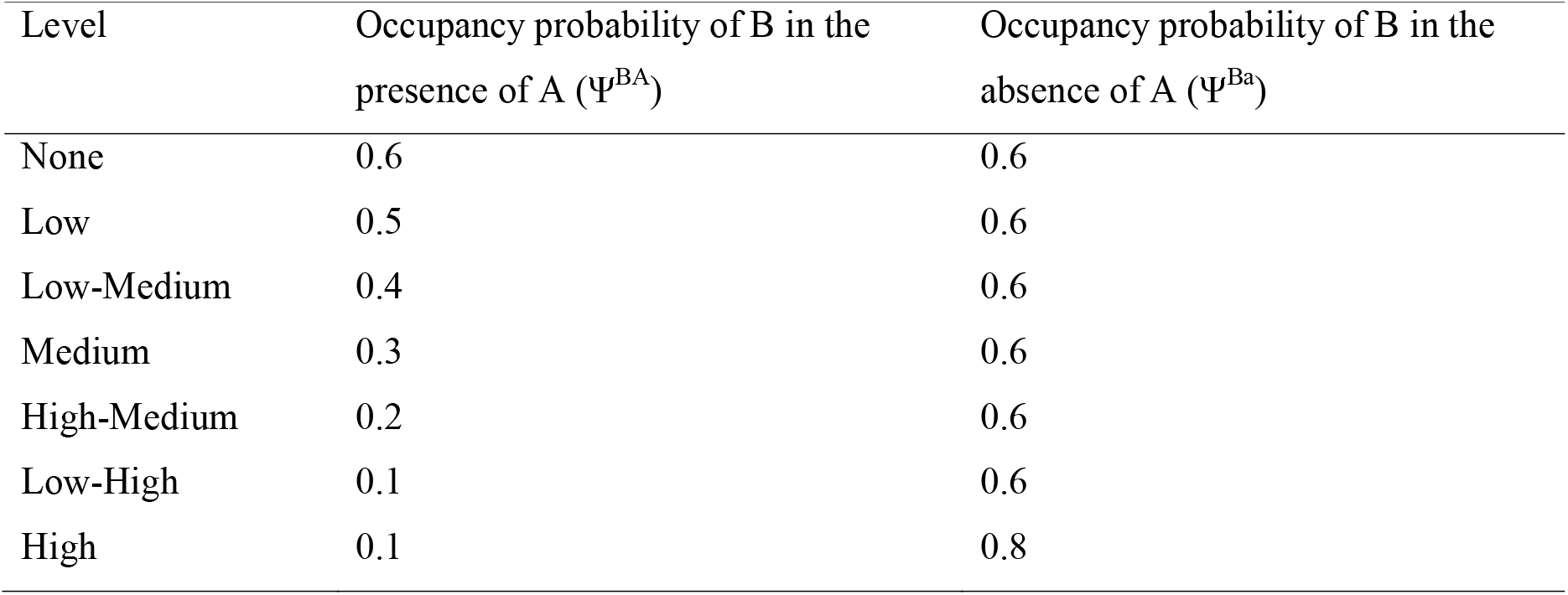
Different scenarios of avoidance used to simulate the presence of B given the presence or absence of A.

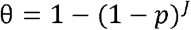

where *J* is the number of temporal replicates.

We set θ^*A*^ to 0.6 and θ^*B*^ to 0.8. We assumed that there was no abundance-induced heterogeneity in θ. To assess the effect of the sampling protocol on the performance of the model, we set the number of temporal replicates to 3; 5 and 10 and set the number of sites to 30; 50; 100; 150; 200 and 250. For each of the resulting 252 scenarios (7 levels of avoidance * 3 levels of temporal replicates * 6 levels of number of sites * 2 values for Ψ^A^), we ran 100 simulations.

To evaluate the performance of the two-species occupancy model under varying levels of avoidance and study designs, we calculated the relative bias and mean squared error (MSE) for Ψ^A^, Ψ^BA^ and Ψ^BA^ by the two-species occupancy model. In a second step, to assess the statistical power to detect the effect of species interactions by the model, we fitted a null model that assumed species were independently distributed and compared the AIC scores of both models. The simulations were done without environmental variation to set a baseline understanding of the study design required to fitted the simplest model.

To evaluate the statistical power of the Rota model (i.e. the capacity of the model to detect interaction when there is interaction), we looked at several metrics:

i. The proportion of simulations in which the model accounting for species interacting performed better than the null model (i.e. had a difference of AIC higher than 2). In the case of ΔAIC > 2, we considered that the co-occurrence was significant.
ii. The proportion of simulations for which the interaction parameter was estimated significant (i.e. had a p-value below 0.05 or with a confidence interval that did not overlap zero).
iii. The proportion of simulations for which the true value of the occupancy parameters was found in the estimated respective confidence intervals (i.e. coverage).

To get an understanding of the precision of the estimated parameters, we also measured the coefficient of variation. The models were fitted using the R package unmarked (Fiske & Chandler, 2011; Kellner et al., 2023).

## Results

### Parameter estimates

#### Mean squared error (MSE)

The MSE improved with increasing level of avoidance and number of sites (Fig.1) irrespective of the values of Ψ^A^ (Appendix 1) or the number of temporal replicates (Appendix 2). Overall, the MSE was the highest for Ψ^BA^ over all the scenarios. These results follow the same trend when Ψ^A^ is set to 0.7 with a difference that the MSE appeared even higher for Ψ^Ba^. The MSE was at its lowest when the avoidance level was at its highest (i.e. “High” scenario) and at its highest in the scenarios in which there was no to very low avoidance. When looking at the sampling protocol, if the number of sites increased, the MSE decreased with the lowest values of MSE for 250 sites, and the highest values of MSE with 30 sites. (Fig. 1). Overall MSE appeared to reach low values once the number of sites was 150 or above.

**Figure 1:**
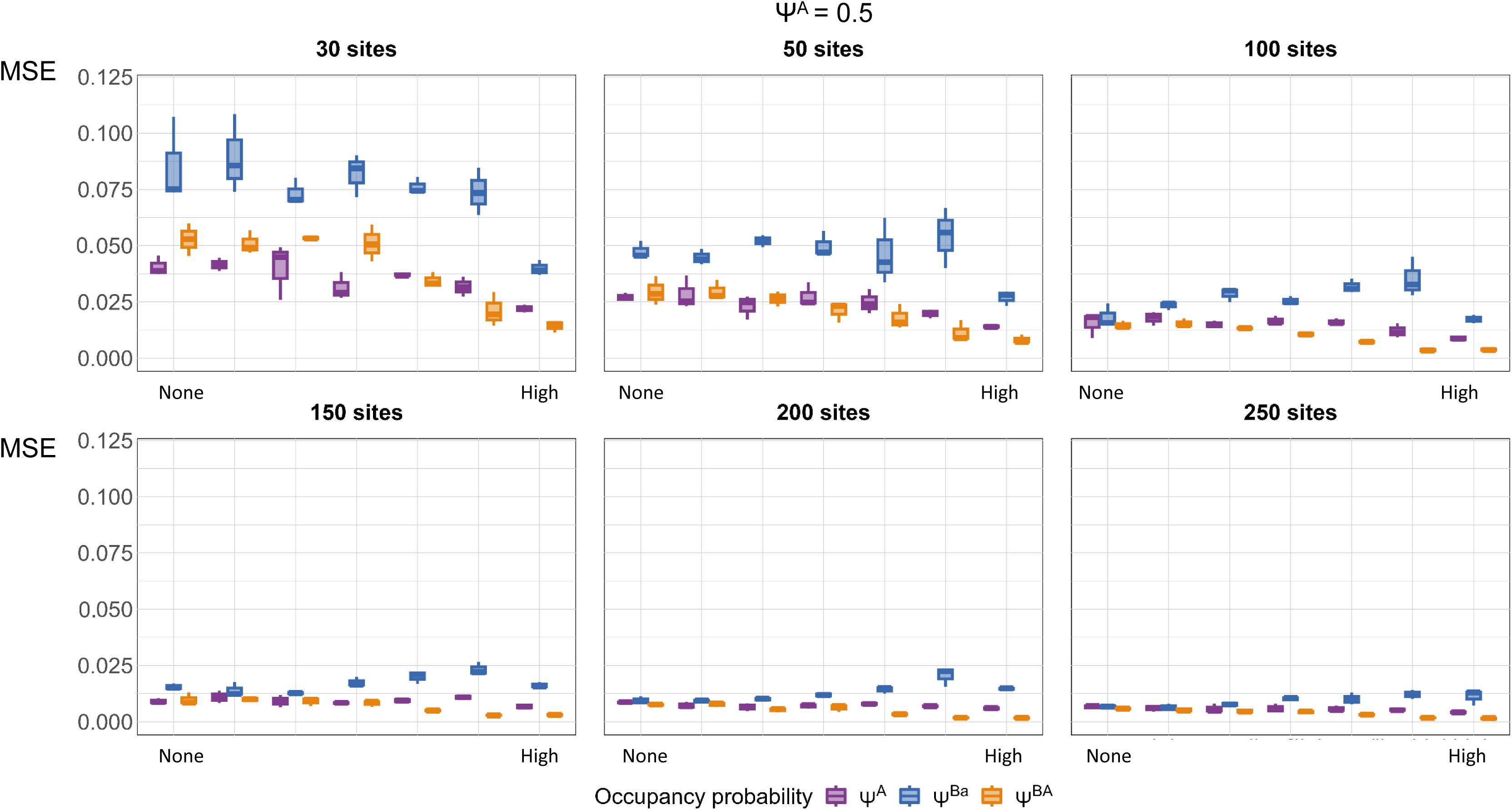
Mean squared errors (MSE) of the different occupancy parameters from our simulation study, and estimated with a 2-species occupancy model by Rota et al. (2016). On the x-axis the MSE was calculated according to different scenarios of avoidance, increasing from no avoidance to high avoidance.

#### Relative bias

The relative bias appeared to decrease with the number of sites (Fig. 2) with all levels of relative bias below 0.05 when the number of sites was at least 150. We found no other clear pattern in the relative bias according to the parameters estimated, the values of Ψ^A^ or the number of temporal replicates (Appendix 1 and Appendix 2). It is worth noting however that Ψ^BA^ was the parameter that often had the highest relative bias.

**Figure 2:**
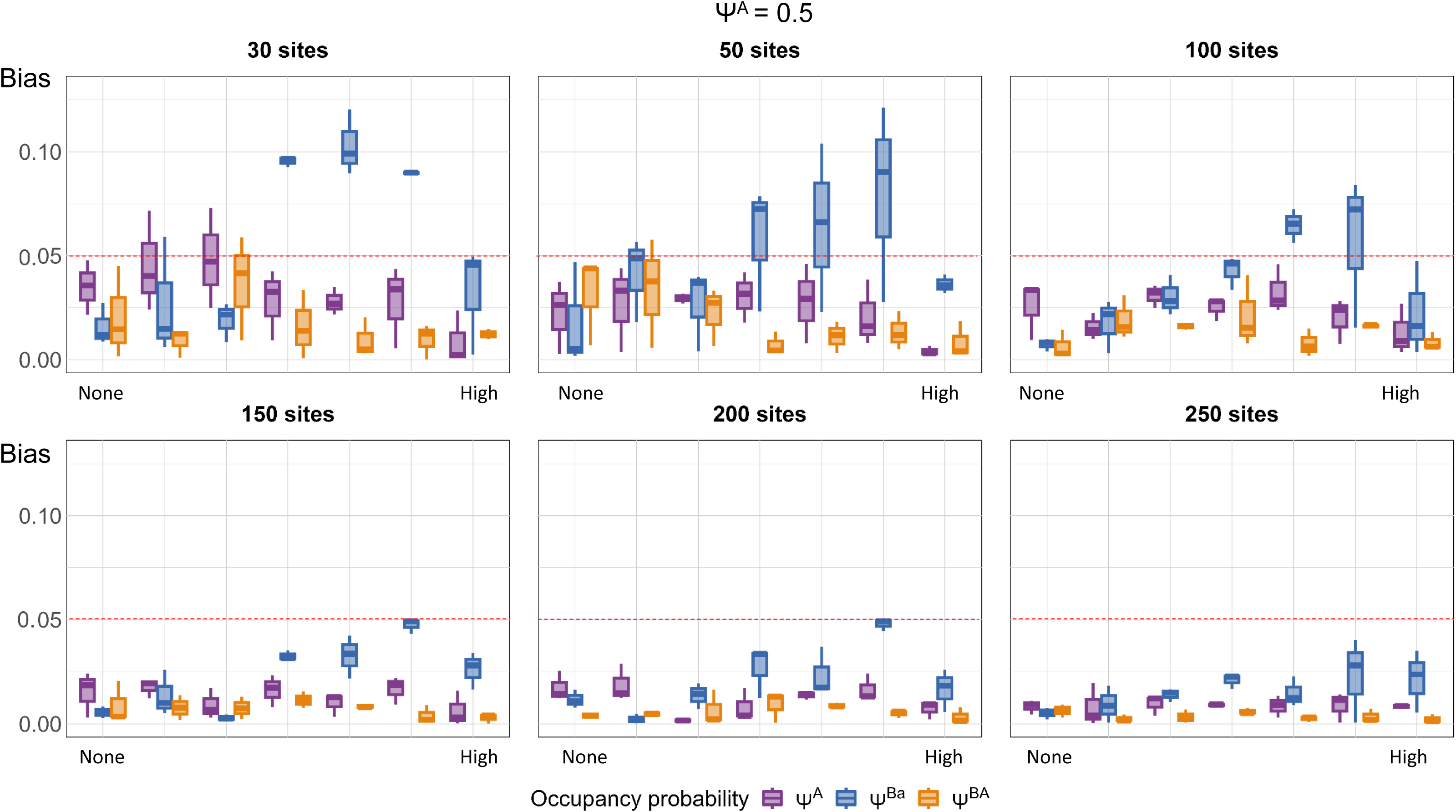
Relative bias of the different occupancy parameters from our simulation study, and estimated with a 2-species occupancy model by Rota et al. 2016. On the x-axis the Relative bias was calculated according to different scenarios of avoidance, increasing from no avoidance to high avoidance.

#### Coefficient of variation

The coefficient of variation appeared to improve when the level of avoidance increased as well as with the number of sites sampled (Fig. 3). For example, we observed high variation (∼50%) when 30 sites were sampled and roughly half as much variation when 250 sites were sampled (Fig. 3). However, the coefficient of variation had a non-linear relationship with respect to interaction strength between species and the number of sites sampled such that it was highest when there were strong interspecific interactions and the number of sites was between 30 and 100 (Fig. 3). The number of repetitions did not seem to have an influence (Appendix 2) and the trend was similar when Ψ^A^ was 0.7 (Appendix 1).

**Figure 3:**
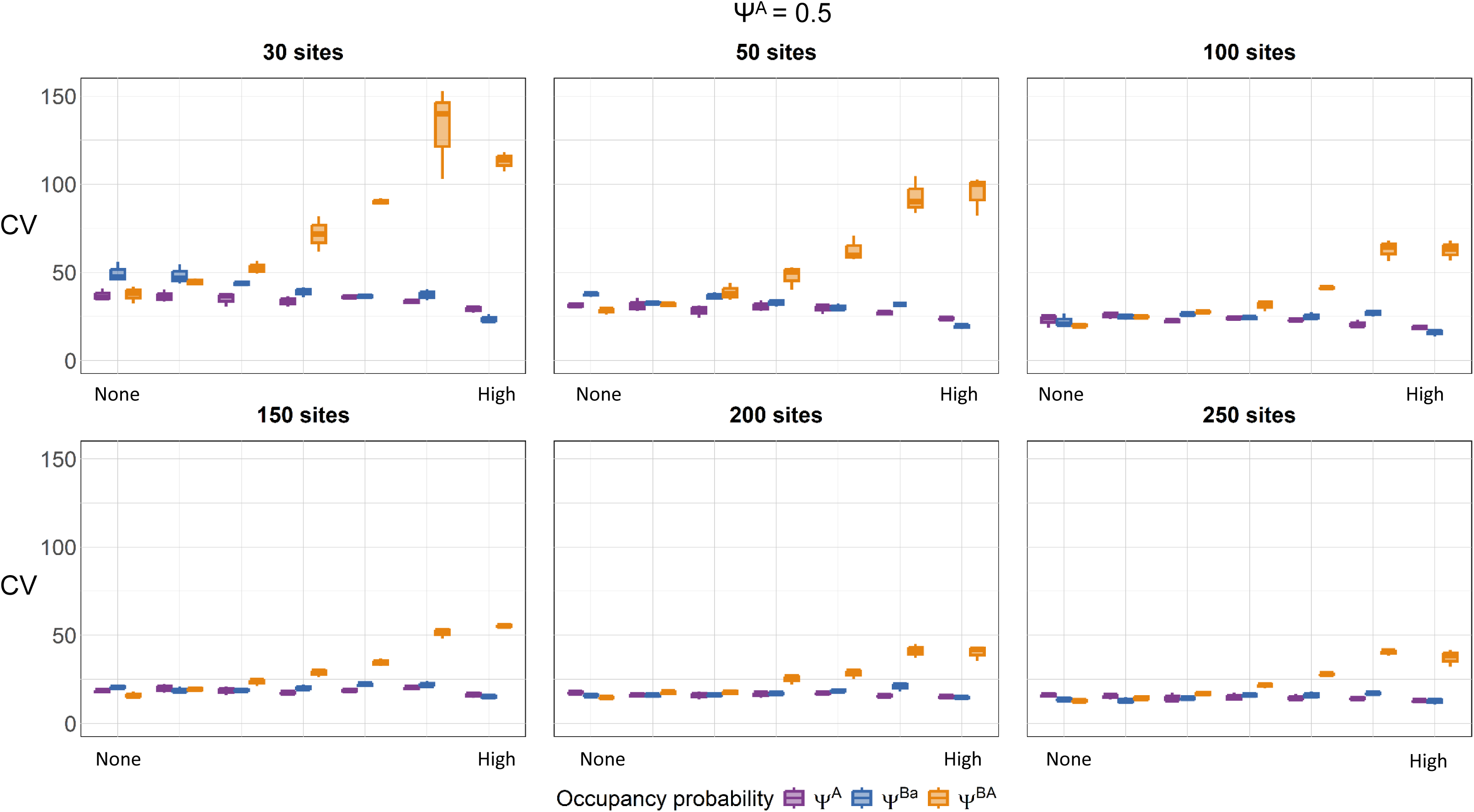
Coefficient of variation of the different occupancy parameters from our simulation study, and estimated with a 2-species occupancy model by Rota et al. 2016. On the x-axis the Relative bias was calculated according to different scenarios of avoidance, increasing from no avoidance to high avoidance.

### Statistical power

The avoidance scenario, the number of sites, and Ψ^A^ were the factors that most influenced the co-occurrence model’s capacity to outperform the occupancy model without species interactions (Fig. 4; left panel, Appendix 1; left panel). When Ψ^A^ was 0.5, to have at least 75% of the simulations having a ΔAIC > 2, the number of sites needed to be higher than 30 and avoidance high. In general, we found that with high levels of avoidance we needed to sample at least 150 sites for the co-occurrence model to have a ΔAIC > 2. Conversely, when there were weak interactions between species the co-occurrence model did not have a ΔAIC > 2 for most simulations across the entire range of sites sampled. However, if Ψ^A^ is 0.7, we found that sampling 200 sites was adequate to have a good chance to detect an interaction between species (Appendix 1). The number of temporal replicates did not influence the capacity of the model to perform better than on occupancy model that would not account for interactions (Fig. 4). When looking at the proportion of simulations that led to a p-value of the interaction parameter below 0.05, it appeared that at least 200 sites needed to be sampled when Ψ^A^ was 0.5. When Ψ^A^ was 0.7, at least 250 sites needed to be sampled. Finally, when looking at the proportion of simulations that covered the true values of occupancy parameters, no clear trend showed as most of the coverage was between 0.85 and 1 (Appendix 3). This is due to the broad confidence intervals that were estimated by the model.

**Figure 4:**
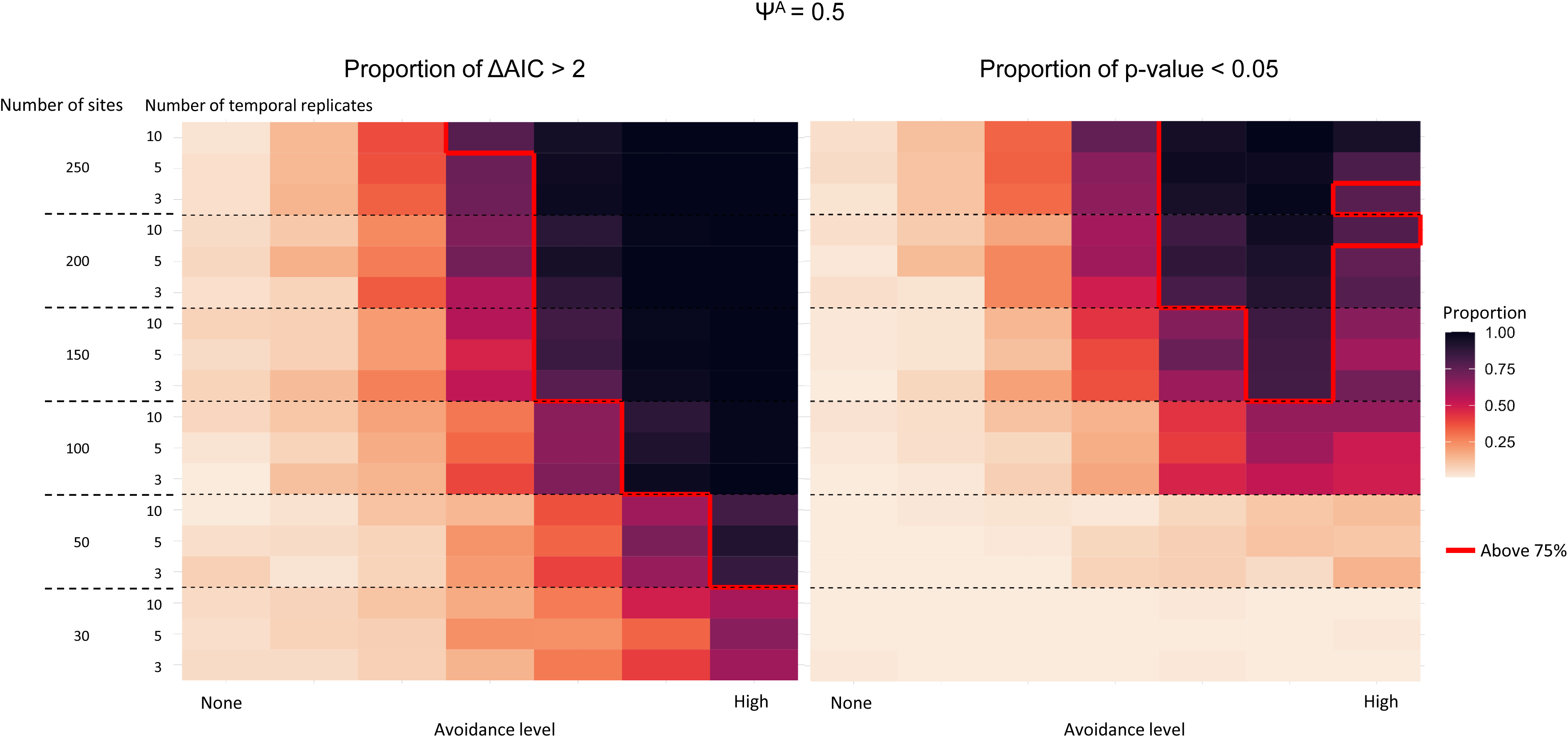
Left panel: Proportion of simulations in which the model accounting for 2 species interacting performed better than an occupancy model not accounting for species interactions (i. e. in which the ΔAIC > 2). On the x-axis this proportion was calculated according to different scenarios of avoidance, increasing from no avoidance to high avoidance. Right panel: Proportion of simulations in which the interaction parameter was significant (i.e. with a p-value > 0.05). On the x-axis this proportion was calculated according to different scenarios of avoidance, increasing from no avoidance to high avoidance.

## Discussion

Our simulation study highlights the importance of sampling protocol to confidently assess co-occurrence patterns with the Rota et al. (2016) co-occurrence model. Over all scenarios, the co-occurrence model did not produce biased estimates (i.e. > 0.05) except for Ψ^Ba^. The precision– represented by the MSE and the coefficient of variation–of the three occupancy parameters decreased with avoidance intensity, while the relative bias was not influenced by avoidance intensity. Additionally, precision was influenced by the number of sites, but not by the number of temporal replicates. Overall, the model produced less precise and more biased results when the true occupancy of the dominant species was higher. Finally, the co-occurrence model only performed better than a model without interactions (i.e. ΔAIC >2 or p-value > 0.05) when avoidance was strong and the number of sites was least 100 to 150. Based on these results, we suggest a minimum of 150 sites when using the Rota et al. (2016) co-occurrence model.

To investigate the statistical power of the Rota et al. (2016) co-occurrence model we made our simulations as simple as possible. We only had two species in our simulations who were neither rare nor exceedingly common (Mackenzie & Royle, 2005; Tingley et al., 2020). Further, each species had a relatively high overall detection probability (MacKenzie et al., 2002; Stauffer et al., 2002) and there was no spatial heterogeneity in occupancy or detection. As such, our results set a baseline suggestion for the minimum number of sites that should be sampled. We expect that if we were to simulate increasingly complex models (e.g. more than two interacting species, accounting for dynamic distribution or environmental covariates) and fit increasingly complex models without a sufficient sample size, and in comparison with our simple framework, the precision would decrease, bias would increase, statistical power would not be improved (Meynard & Kaplan 2012; Meynard et al. 2019), and that the minimum number of sites to sample would have to be larger than 150. Such a study has been recently conducted to assess the performance of the Rota et al. (2016) co-occurrence model in which they compared the simple model (i.e. only two species interacting) to more complex models, with more than two species and increasing number of environmental covariates (Cowans et al., 2024). Their simulations reveal high bias and low coverage in co-occurrence estimates with fewer than 400 sites, though strong interactions become reliable above this threshold. Weak interactions are undetectable even with large samples suggesting caution when applying these models to small datasets or when interactions are weak.

As expected, the precision and statistical power of the model we tested were influenced by avoidance intensity, with these two measures improving when the avoidance (i.e., effect size) increased. However, the number of temporal replicates did not affect this relationship. Given that we set the detection probability quite high, there were very few cases where false negatives occurred on sites throughout the repetitions. It is worth noting that relative bias did not seem to be influenced by the avoidance scenario or sampling protocol. Overall, the relative bias remained under 0.05 which we considered good. To further investigate such performance, we suggest exploring a Bayesian approach (e.g. Green et al. (2018); Fidino et al. (2019)) to deal with the issue of boundary estimates that is known to influence precision in single-species model parameter estimation (see Table 10.2 in Kéry & Royle 2021).

When comparing models via AIC or p-value of the interaction parameter, we found that the models were equivalent across most scenarios both models. This means that, for low to mild levels of avoidance, the co-occurrence model was not the most supported one even though the data generating parameters assumed interactions were present. These scenarios are an example of the many cases where a non-significant result occurs, but the interaction might not be strong enough for the null hypothesis to be rejected. We emphasize that the absence of evidence for an interaction does not necessarily imply that no interaction exists.

When comparing our results with the 13 empirical studies that used the Rota et al. (2016) framework, 4 studies had less than 150 sites (Fox-Rosales & de Oliveira, 2023; Gámez & Harris, 2021; Lahkar et al., 2021; Saeed et al., 2022) but all 4 studies found a significant interaction parameter. This highlights the need to further include ecological complexity, such as environmental gradient on species’ presence and interactions when simulating and assessing species interactions. For studies with elusive species, we encourage the use of simulations to determine an adequate sample size, which could be informed by data. Co-occurrence strength considerably varied across studies that have used the Rota et al. (2016) framework. For example, avoidance strength has varied from low (e.g. Siberian ibex and the grey wolf, Salvatori et al. (2021); black bear and Grizzly bear; Ladle et al. (2018), to high (e.g. grey squirrel and marten in broadleaf environment; Twining et al. (2021), and changes along environmental gradients (e.g. Kass et al. (2020); Parsons et al. (2022). When looking at the sampling protocols that were used in these studies, the number of sites varied greatly (e.g. from 14 sites, Saeed et al. (2022); to 1427 sites, Parsons et al. (2018)) as well as the number of temporal replicates (e.g. 4 temporal replicates, Kass et al. (2020); to more than 20 temporal replicates, Ladle et al. (2018)). Of the 13 empirical studies that used the Rota et al. (2016) co-occurrence framework, most reported p-values or confidence intervals for their interaction parameter while only 2 compared their model with another that assumed species were independently distributed. In both of these papers, the co-occurrence model outperformed the simpler parameterization (Fox-Rosales & de Oliveira, 2023; Saeed et al., 2022).

We acknowledge that increasing the number of sampling units can present some logistical constraints. One solution to this problem is to collect data at sites over multiple sampling seasons. Such a study design makes it possible to use an auto-logistic parameterization of this co-occurrence model (Zipkin et al., 2012), such as in Kass et al. (2020), which used a first-order auto-logistic term that accounts for changes in the occupancy status of a species if they were present in the preceding time-step. While collecting data over time also implies that a dynamic formulation could be used (e.g., Fidino et al. (2019)), the auto-logistic parameterization introduces far fewer parameters, making it appealing to studies with a more limited sample size. Alternatively, integrated models present a novel framework for interacting species in which data from several difference sources that vary in information content can be integrated and unified (Barraquand & Gimenez, 2019; Quéroué et al., 2021). Such models have proven to improve the accuracy and precision of the parameters of interest and represent promising solutions (Isaac et al., 2020; Zipkin et al., 2023).

In our study, we assessed patterns of co-occurrence between species. However, whether species interactions are reflected by co-occurrence patterns remains unclear (Blanchet et al., 2020). Indeed, the presence of multiple species within the same habitat does not necessarily imply direct or indirect interactions. Co-occurrences can be a result of shared habitat preferences, interactions with other species that are not accounted for, similar environmental tolerances, or stochastic processes that need to be integrated in complex models (Dormann et al., 2018). Yet, even if co-occurrence may not provide evidence of ecological interactions, understanding these patterns is still important.

Overall, our study highlights the need to further develop methods that allow detecting the signal of interactions even in low to mild interaction levels with a low number of sites. The number of statistical tools we have available now is much greater than it was in the past. However, to appropriately use these models we need to know what appropriate sample sizes are. Here, we show that in the simplest cases, at least 150 sites are necessary. We do not argue that this is the minimum required for a study to be published with this model, especially as our simulations will not capture the range of possible patterns that could be captured in nature, but simulations, informed by data could provide information into the possible range of sites that are required.

## Data accessibility

The scripts to run, analyse the simulations and plot the results can be found in the following GitHub repository: https://github.com/JulieLouvrier/RotaSimulStudy_repo (DOI: 10.5281/zenodo.14007815)

## Author contributions

**Julie Louvrier:** Conceptualization, Methodology, Writing - original draft. **Olivier Gimenez:** Conceptualization, Writing - review & editing, Supervision. **Mason Fidino:** Conceptualization, Writing - review & editing. Josefa Vergara Stuardo: Writing - review & editing.

## Acknowledgements

The project was funded through a National Research Agency grant DEMOCOM ANR-16-CE02-0007 (Effets de la gestion et du climat sur la dynamique des communautés – Développement d’une démographie multi-espèces).

