## Supplementary figures and images for "How reliably can species co-occurrence be inferred from occupancy models?"

### Appendix 1

#### Appendix 1: ΔAIC, Bias, MSE and CV for psiA = 0.7

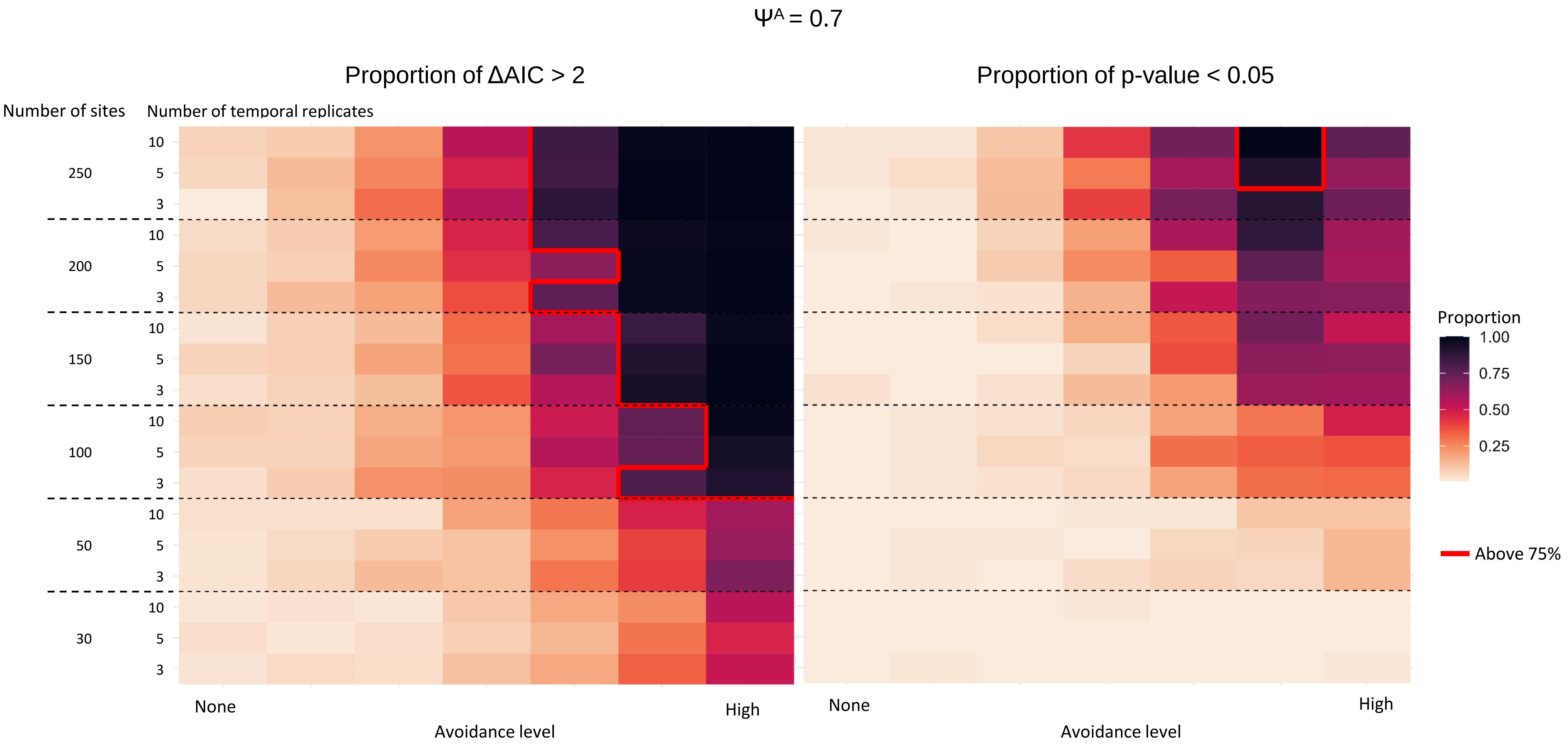

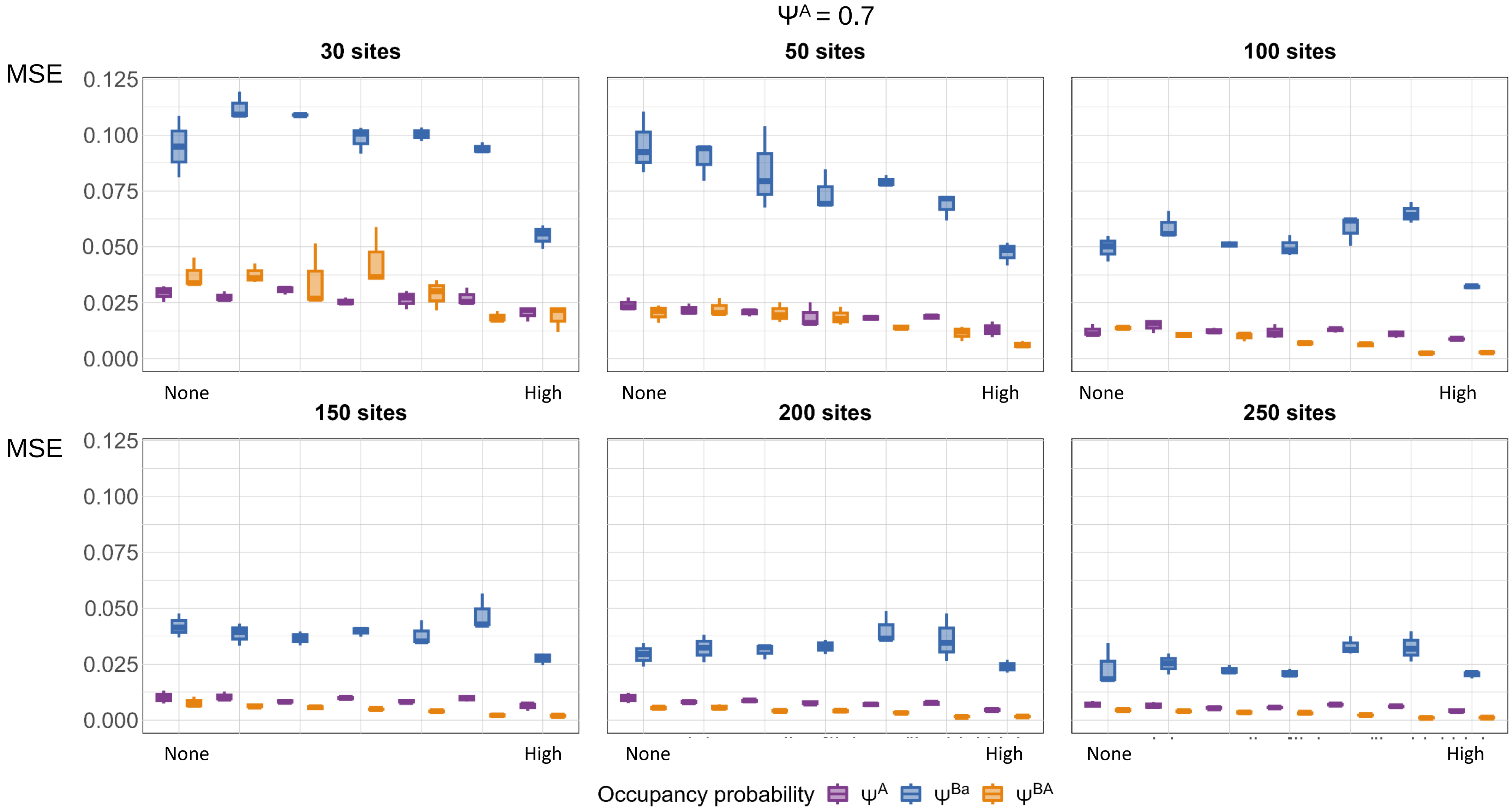

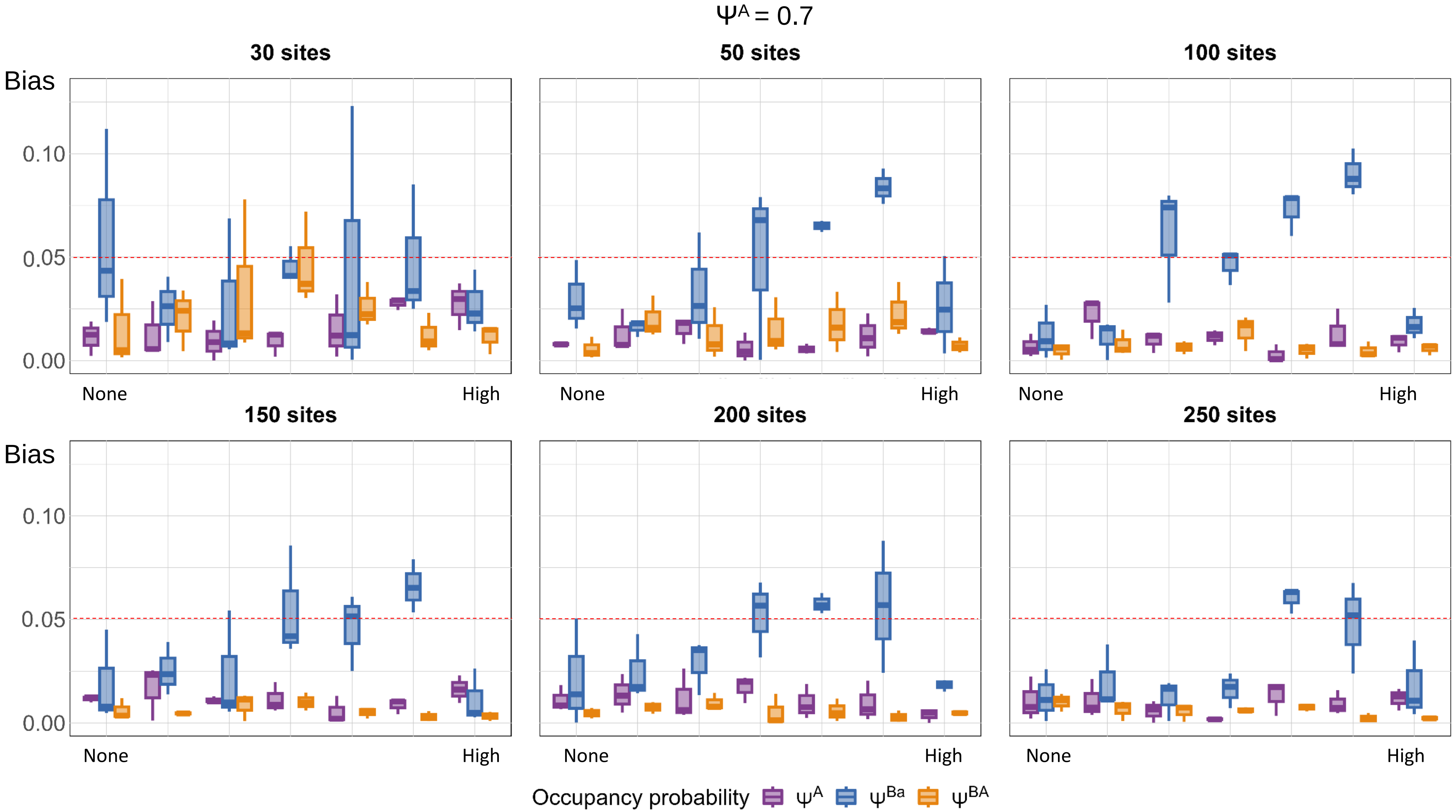

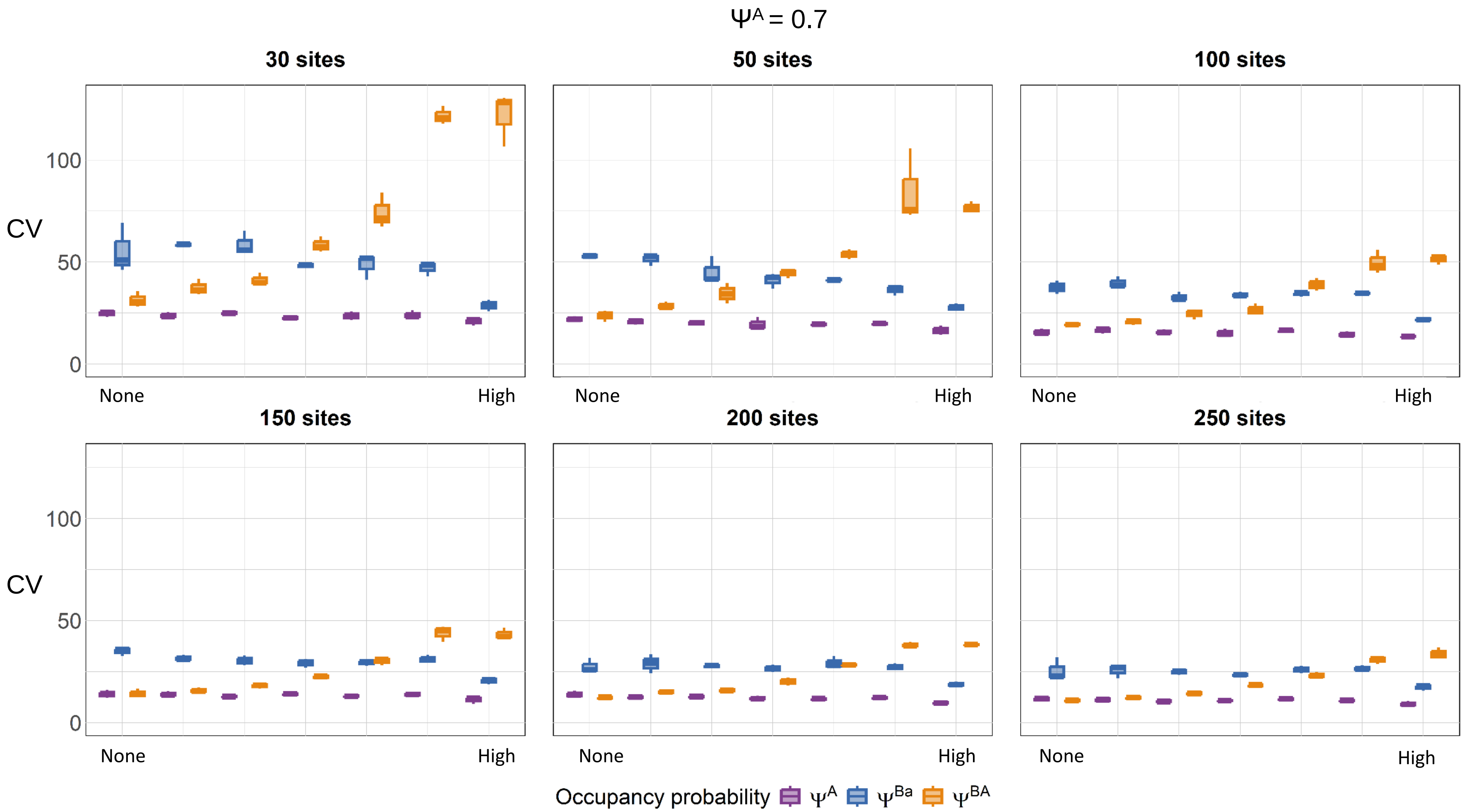

### Appendix 2

#### Appendix 2: Bias, MSE and CV depending on the number of repetitions

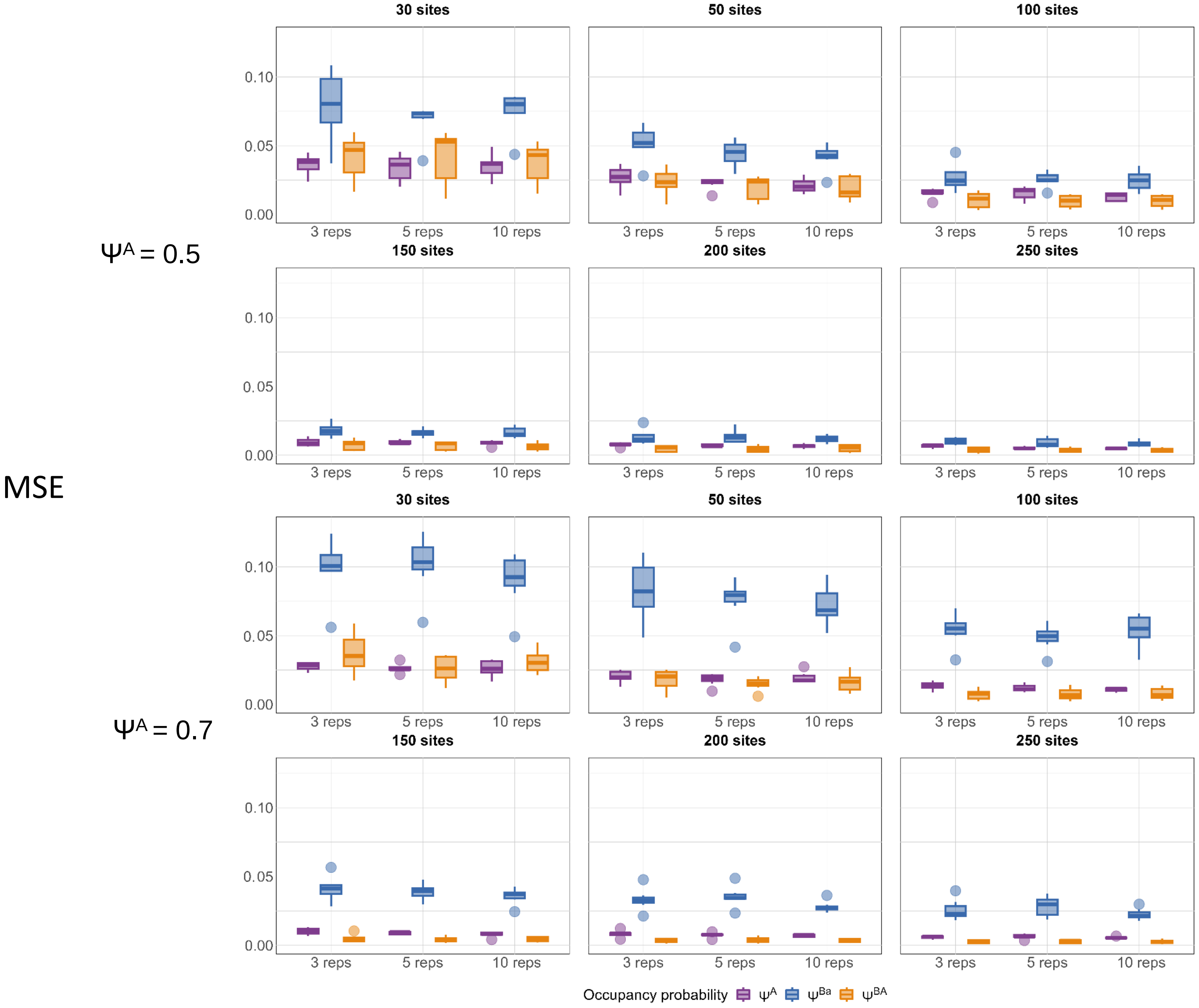

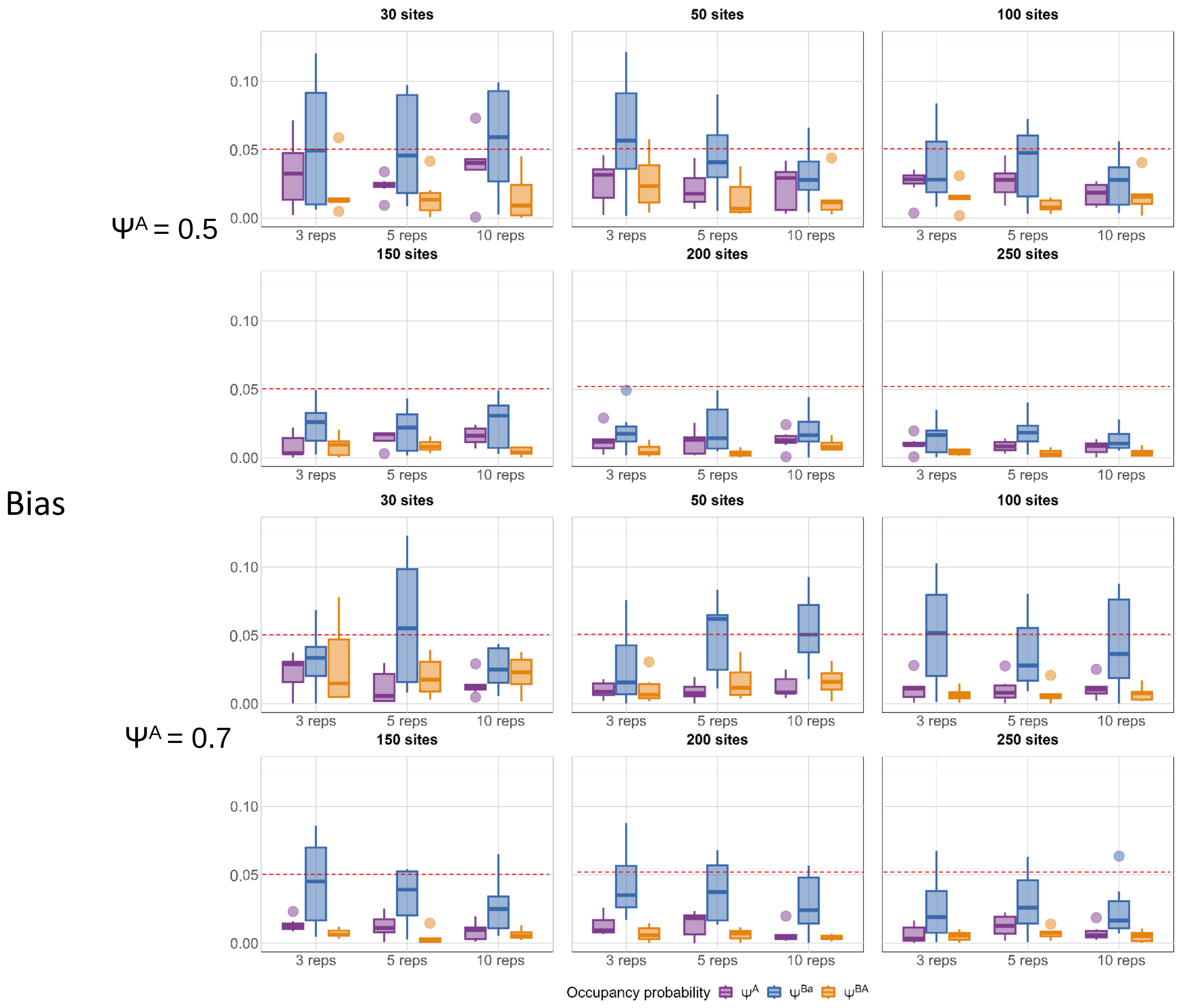

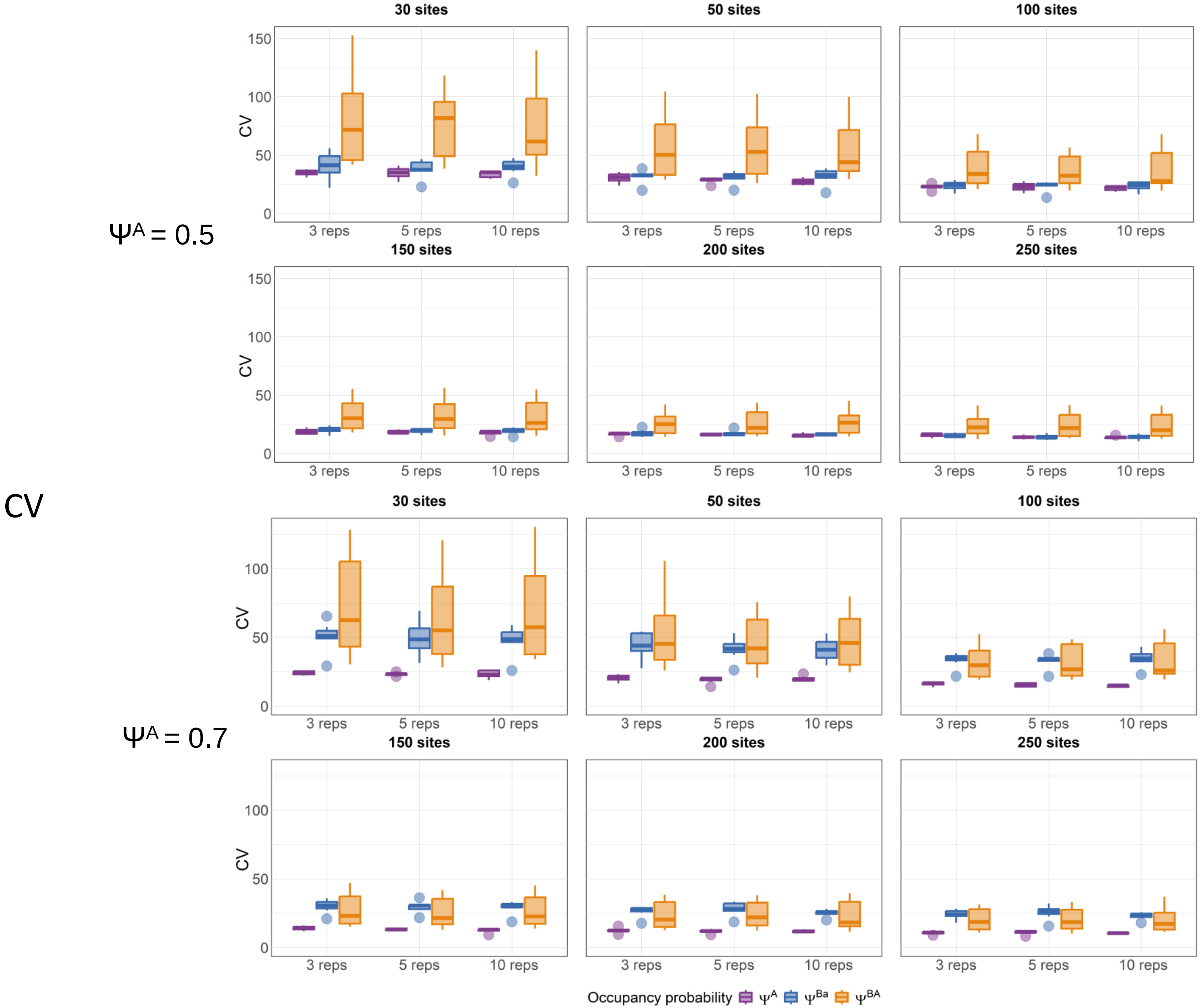

### Appendix 3

#### Appendix 3 : Coverage

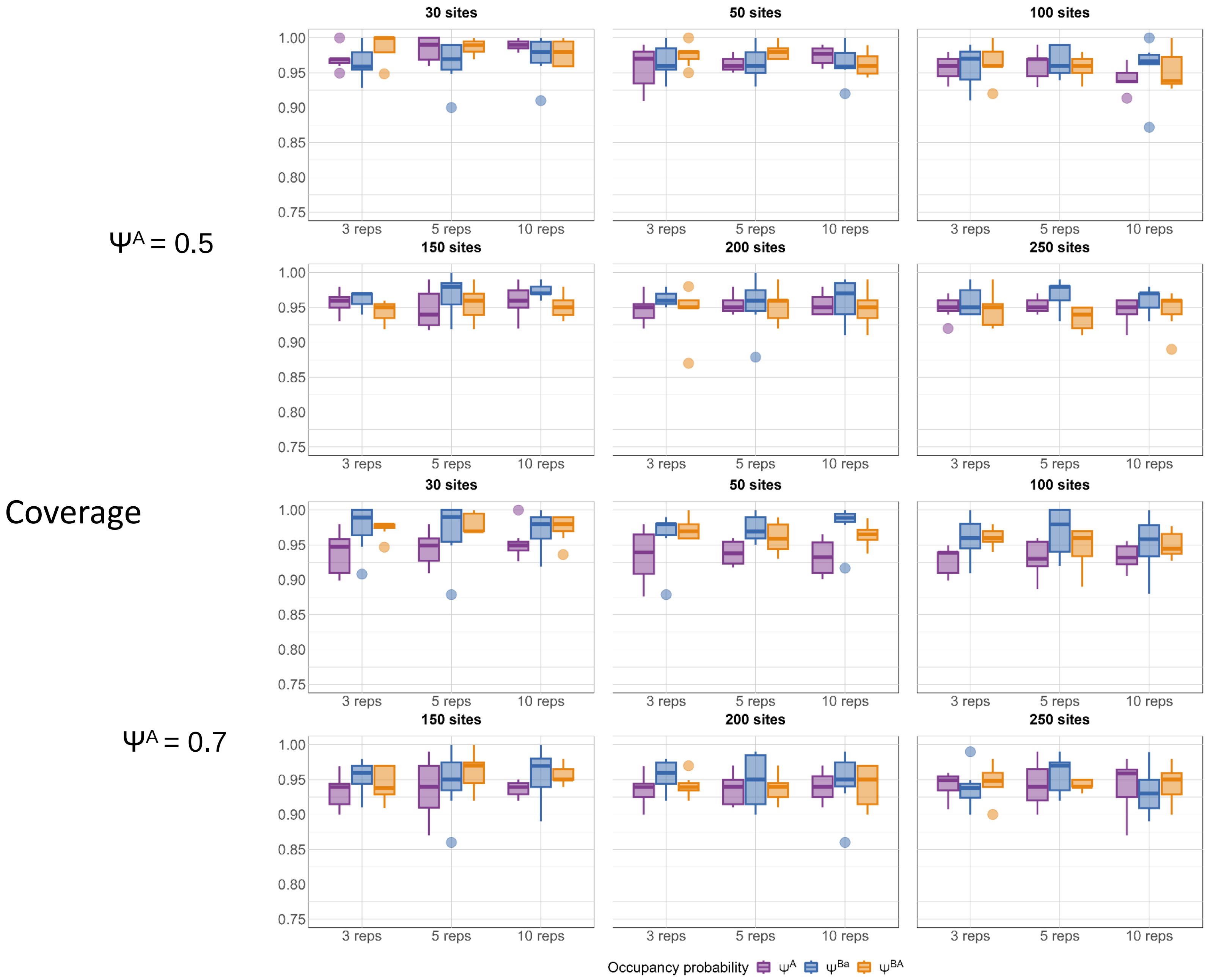

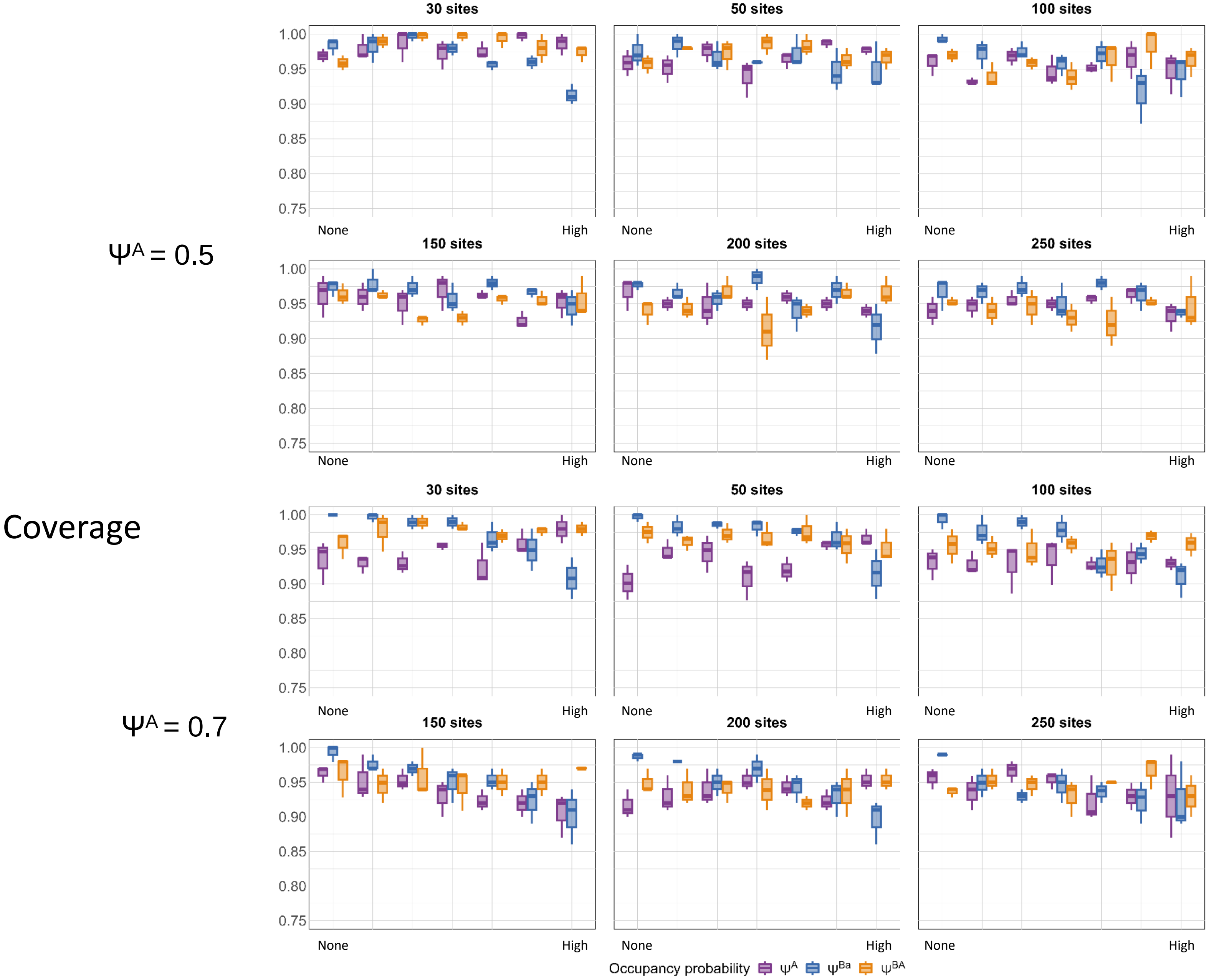
